# NPAS4 in the medial prefrontal cortex mediates chronic social defeat stress-induced anhedonia and dendritic spine loss

**DOI:** 10.1101/2021.03.04.433930

**Authors:** Brandon W Hughes, Benjamin M Siemsen, Stefano Berto, Jaswinder Kumar, Rebecca G Cornbrooks, Rose Marie Akiki, Jordan S Carter, Michael D Scofield, Christopher W Cowan, Makoto Taniguchi

## Abstract

Chronic stress can produce reward system deficits (*i*.*e*. anhedonia) and other common symptoms associated with depressive disorders, as well as neural circuit hypofunction in the medial prefrontal cortex (mPFC). However, the molecular mechanisms by which chronic stress promotes depressive-like behavior and hypofrontality remain unclear. We show here that the neuronal activity-regulated transcription factor, NPAS4, in the mPFC is regulated by chronic social defeat stress (CSDS), and it’s required in this brain region for CSDS-induced changes in sucrose preference and natural reward motivation. Interestingly, NPAS4 is not required for CSDS-induced social avoidance or anxiety-like behavior. We also find that mPFC NPAS4 is required for CSDS-induced reduction of pyramidal cell dendritic spine density, revealing a relationship between mPFC dendritic spine changes and anhedonia-like behavior, but not social avoidance behavior. Finally, transcriptomic analysis from the mPFC revealed that NPAS4 influences expression of numerous genes linked to glutamatergic synapses and ribosomal function, as well as many dysregulated genes observed in common neuropsychiatric disorders, including depression. Together our findings reveal an essential role for the activity-regulated transcription factor, NPAS4, in chronic stress-induced mPFC hypofunction and anhedonia.

## Introduction

Stress-related mental disorders continue to be a leading cause of disability and financial burden on society ^1^. The associated symptom domains of stress-related disorders are diverse and present with a high degree of comorbidity, thus treatment strategies for these disorders represent a major healthcare challenge. The rodent chronic social defeat stress (CSDS) paradigm produces multiple behavioral and neural phenotypes reminiscent of stress-related and depressive disorders in humans, including anhedonia-like behaviors and social avoidance ^2-11^. CSDS produces social avoidance in some, but not all, mice, which is similar to human responses to chronic stress ^6, 9, 12^. Another CSDS-induced behavior is anhedonia, a core symptom of major depressive disorder (MDD) that is associated with deficits in hedonic capacity, reward evaluation, decision-making, and motivation to obtain rewards, as well as risk for suicide and treatment resistance ^13-17^. However, the neural mechanisms by which chronic stress produces anhedonia remain unclear. Multiple preclinical and clinical studies have revealed reduced function of the medial prefrontal cortex (mPFC), which is caused, at least in part, by stress-induced loss of structural and functional synaptic connections and neural circuits in this region ^3, 18-21^. Furthermore, this stress-induced hypofrontality is thought to underlie symptoms of MDD ^14, 22-24^ and contribute to the neuropathology of treatment-resistant depression ^25^, including the potential for anhedonia susceptibility ^26, 27^.

In this study, we investigated the role of Neuronal PAS domain Protein 4 (NPAS4) in chronic stress-induced brain and behavior dysfunction. NPAS4 is an immediate early gene (IEG) and transcription factor that modulates synaptic connections on excitatory (E) and inhibitory (I) neurons in response to synaptic activity – a proposed homeostatic mechanism to modulate E/I balance in strongly activated neural circuits ^28-34^. In the brain, NPAS4 is required in the hippocampus and amygdala for contextual fear learning ^35, 36^, in the visual cortex for social recognition ^37^, and in the nucleus accumbens (NAc) for cocaine reward-context learning and memory ^38^. As such, NPAS4 is well-positioned to mediate adaptive cellular and synaptic changes produced by strong circuit activity, such as that produced in the mPFC by acute and chronic stress. Here, we discovered that acute and chronic social defeat stress induce NPAS4 expression in the mPFC, and NPAS4 in this brain region is required for both CSDS-induced anhedonia and reduction of pyramidal neuron dendritic spine density. Similarly, we found that reducing mPFC *Npas4* dysregulates numerous downstream genes that are important for ribosome function or excitatory synapse organization, activity, and signaling – the majority of which are differentially expressed in human patients with MDD ^39^. Our findings revealed an essential role for a NPAS4 in chronic stress-induced mPFC hypofrontality and anhedonia.

## Methods and Materials

### Recombinant plasmids and shRNA expression viral vectors

For knockdown of endogenous *Npas4* mRNA expression in mPFC, previously validated *Npas4* shRNA or scramble (SC) shRNA control was cloned into the pAAV-shRNA vector as previously described ^29, 38^. The adeno-associated virus serotype 2 (AAV2) vector consists of a CMV promoter driving eGFP with a SV40 polyadenylation signal, followed downstream by a U6 RNA polymerase III promoter and *Npas4* shRNA or scrambled (SC) shRNA oligonucleotides, then a polymerase III termination signal - all flanked by AAV2 inverted terminal repeats. AAV2-*Npas4* shRNA and SC shRNA were processed for packaging and purification by the UNC Vector Core (Chapel Hill, NC).

### Viral-mediated gene transfer

Stereotaxic surgery was performed under general anesthesia with a ketamine/xylazine cocktail (120 mg/kg: 16 mg/kg) or isoflurane (induction 4% v/v, maintenance 1%–2% v/v). Coordinates to target the mPFC (ventral portion of cingulate, prelimbic, and infralimbic cortices) were +1.85-1.95 mm anterior, +0.75 mm lateral, and 2.65 to 2.25 mm ventral from bregma (relative to skull) at a 15° angle in all mice ^3^. AAV2-scramble (SC) shRNA (2.9*10^^9^ and 1.1 *10^12^ GC/mL) and AAV2-*Npas4* shRNA (4.3*10^^9^ and 3.1*10^12^ GC/mL) were delivered using Hamilton syringes or nanoinjectors with pulled glass capillaries at a rate of 0.1 μL/min for 0.4 μL total at the dorsoventral sites, followed by raising the needle and an additional 0.4 μL delivery of virus. After waiting for an additional 5-10 min, needles were completely retracted. Viral placements were confirmed through immunohistochemistry for bicistronic expression of eGFP from the AAV2 viral vectors by experimenters blinded to the experimental conditions. Animals with off-target virus infection or no infection in one or both hemispheres were excluded from the analysis of behavioral phenotypes.

### Chronic social defeat stress

Chronic social defeat stress (CSDS) was performed as previously described ^5, 6^. CD1 retired male breeders (Charles River Laboratory, CA) were single-housed for 3-5 days before CSDS procedures to establish their territorial cage, then pre-screened for aggressive behavior. Experimental C57BL/6J male mice were introduced to the aggressor’s territorial cage, physically contacted and attacked by the aggressor for 5-10 min, and then separated by a clear plastic board with multiple small holes for 24 hours. Experimental mice were introduced to a new CD1 aggressor each day. The no stress control mice were housed with another non-stressed C57BL/6J male mouse, separated by the same plastic board, and the cage partner was changed every day for 10 days of the CSDS experiment.

## Results

### Social defeat stress induces NPAS4 expression in the medial prefrontal cortex

We first examined the expression of *Npas4* mRNA in two key corticolimbic regions associated with stress and reward, the mPFC and the nucleus accumbens (NAc), following 11 days of CSDS. We compared CSDS responses to a single social defeat stress experience (acute stress; Figure 1A). In the mPFC, we observed a very rapid and transient induction of *Npas4* mRNA in the mPFC (Figure 1B) and NAc (Supplemental Figure S1A). We observed a similar response with *cFos* mRNA, albeit a slower induction and longer duration (Supplemental Figure S1B). Interestingly, CSDS-induced expression of both *Npas4* and *cFos* was observed, although it was reduced compared to the acute stress response (Figure 1B; Supplemental Figure S1B), possibly due to CSDS-induced mPFC hypofunction. In contrast, the CSDS-induced attenuation of *Npas4* induction was not observed in the NAc (Supplemental Figure S1A). Similar to *Npas4* mRNA induction, we observed a significant increase in cells expressing NPAS4 protein at 1 hour following CSDS or acute stress exposure in multiple mPFC regions, including the anterior cingulate and prelimbic cortex subregions (Figure 1C). Interestingly, the vast majority (>75%) of NPAS4+ neurons in the mPFC were CaMKIIα-expressing pyramidal neurons (Figure 1D, 1E), with almost no detectable NPAS4 expression in parvalbumin- or somatostatin-expressing GABAergic interneurons. Similar to *Npas4* mRNA, the relative NPAS4 protein expression per cell was highest following acute stress (Supplemental Figure S1C), suggesting that acute stress and CSDS activate a similar number of NPAS4-positive mPFC neurons, but the NPAS4 expression level within each cell is lower following repeated psychosocial stress (Supplemental Figure S1C).

**Figure 1.**
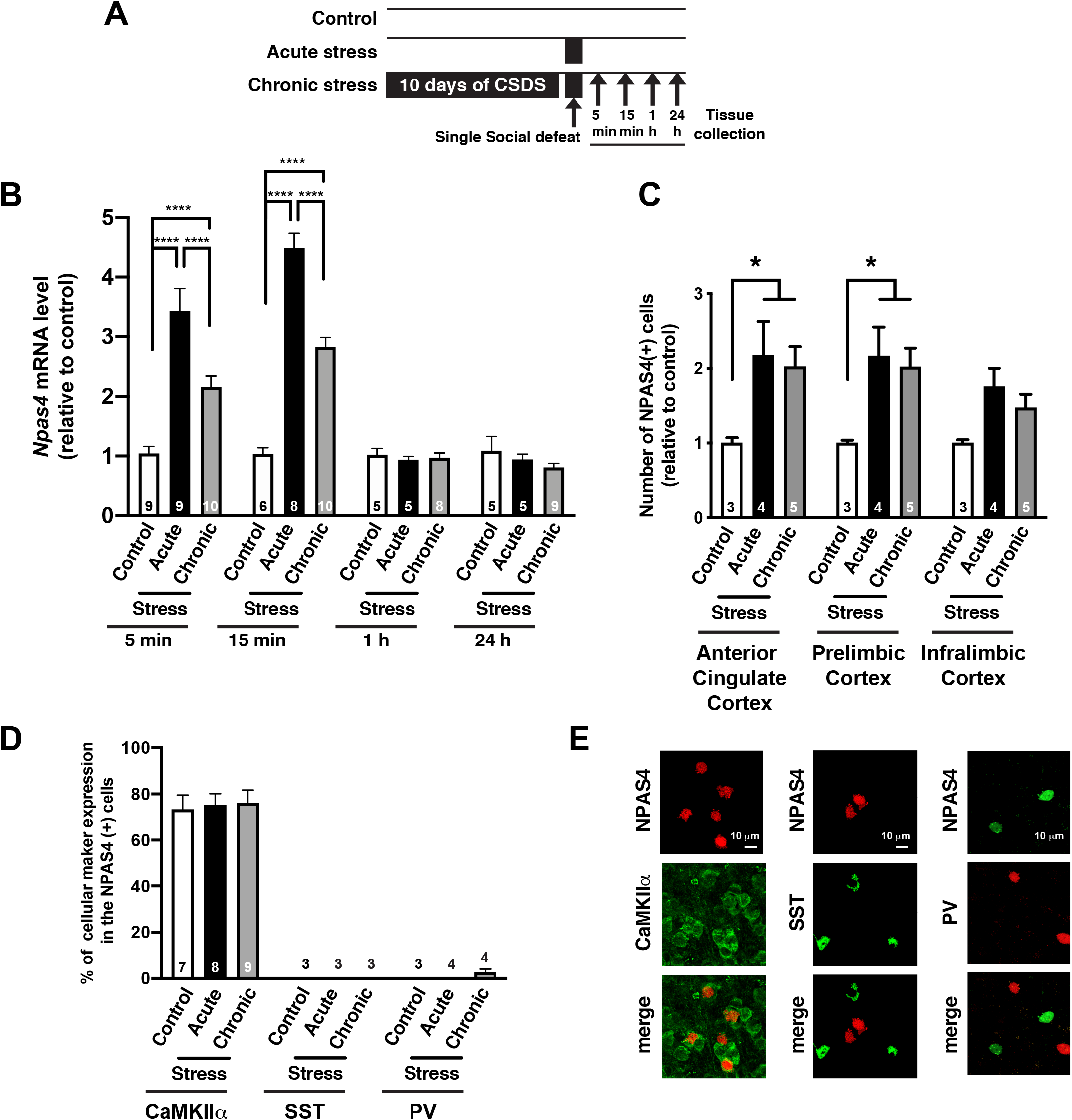
Social defeat stress induces NPAS4 expression in the mPFC. (A) Schematic illustration of experimental timeline of gene expression analyses following acute social defeat stress and 10 days of chronic social defeat stress (CSDS). (B) Data plot represents the quantification of *Npas4* mRNA expression following acute and chronic social defeat stress at 5 min, 15 min, 1h, and 24 hr (n= 5-10/condition). (C) Quantification of fold change in NPAS4-positive cell number following acute and chronic social defeat stress in subregions of the mPFC, including the anterior cingulate, prelimbic, and infralimbic cortices (n= 3-5/condition). (D and E) Data plot shows the percentage of CaMKIIα-, Somatostatin (SST)-, and Parvalbumine (PV)-positive cells in NPAS4-positive cells within the mPFC after acute stress and CSDS (n= 3-9/condition), as well as representative IHC images of NPAS4 colocalization in these respective cell type. Scale bar, 10 μm. Data shown are mean ± SEM; *p < 0.05, ****p < 0.0001. Also see Table S1 for detailed statistical analyses.

### NPAS4 in the mPFC is required for CSDS-induced anhedonia-like behavior

To examine the function of NPAS4 in CSDS-induced behaviors (Figure 2A), we employed a neurotropic AAV-mediated RNA-interference approach to reduce endogenous *Npas4* in the mPFC (AAV2-*Npas4* shRNA^PFC^ and Figure 2B) ^38^. Adult male mice (C57BL/6J) received a bilateral injection of AAV2-*Npas4* shRNA^PFC^ or AAV2-shRNA scrambled control (AAV2-SC shRNA^PFC^). Subsequently, the mice were subjected to 10 days of CSDS or no stress control condition, and then they were tested for sociability, natural reward preference and motivation, and anxiety-like behavior (Figure 2A). The CSDS-treated SC shRNA^PFC^ and *Npas4* shRNA^PFC^ mice showed a significant reduction in the time spent interacting with a novel social target, as shown by time spent in the interaction zone in the presence a social target (Figure 2C). In addition, there was a main effect of CSDS, but no significant difference between *Npas4* shRNA^PFC^ vs. SC shRNA^PFC^ mice in the relative distribution of social interaction ratio in CSDS-treated mice (Figure 2D). Both *Npas4* shRNA^PFC^ and SC shRNA^PFC^ mice showed significantly increased social avoidance time and ratio following CSDS (Figure 2E and 2F), suggesting that mPFC NPAS4 is not required for CSDS-induced social avoidance. However, unlike the CSDS-treated SC shRNA^PFC^ mice, CSDS-treated *Npas4* shRNA^PFC^ mice did not develop anhedonia-like behavior, as detected by a significant reduction in the sucrose preference in the 2-bottle choice test (Figure 2G and supplemental Figure S2A). Interestingly, CSDS increased anxiety-like behavior, as measured in the elevated plus maze, in both SC shRNA^PFC^ and *Npas4* shRNA^PFC^ mice (Figure 2H), indicating that mPFC NPAS4 function is required for some, but not all, of the behavioral sequelae of CSDS. This was further confirmed by correlation analysis, which showed that neither sucrose preference nor decreased anxiety were statistically correlated with the social interaction ratio in either group (Supplemental Figure S2B and S2C), suggesting that the molecular and circuit mechanisms of CSDS-induced social avoidance, anhedonia, and anxiety might be distinct. Moreover, the presence of CSDS-induced social avoidance and anxiety-related behavior in *Npas4* shRNA^PFC^ mice argues against the possibility that they are simply less sensitive to stress and/or have deficits in threat/fear-related learning and memory.

**Figure 2.**
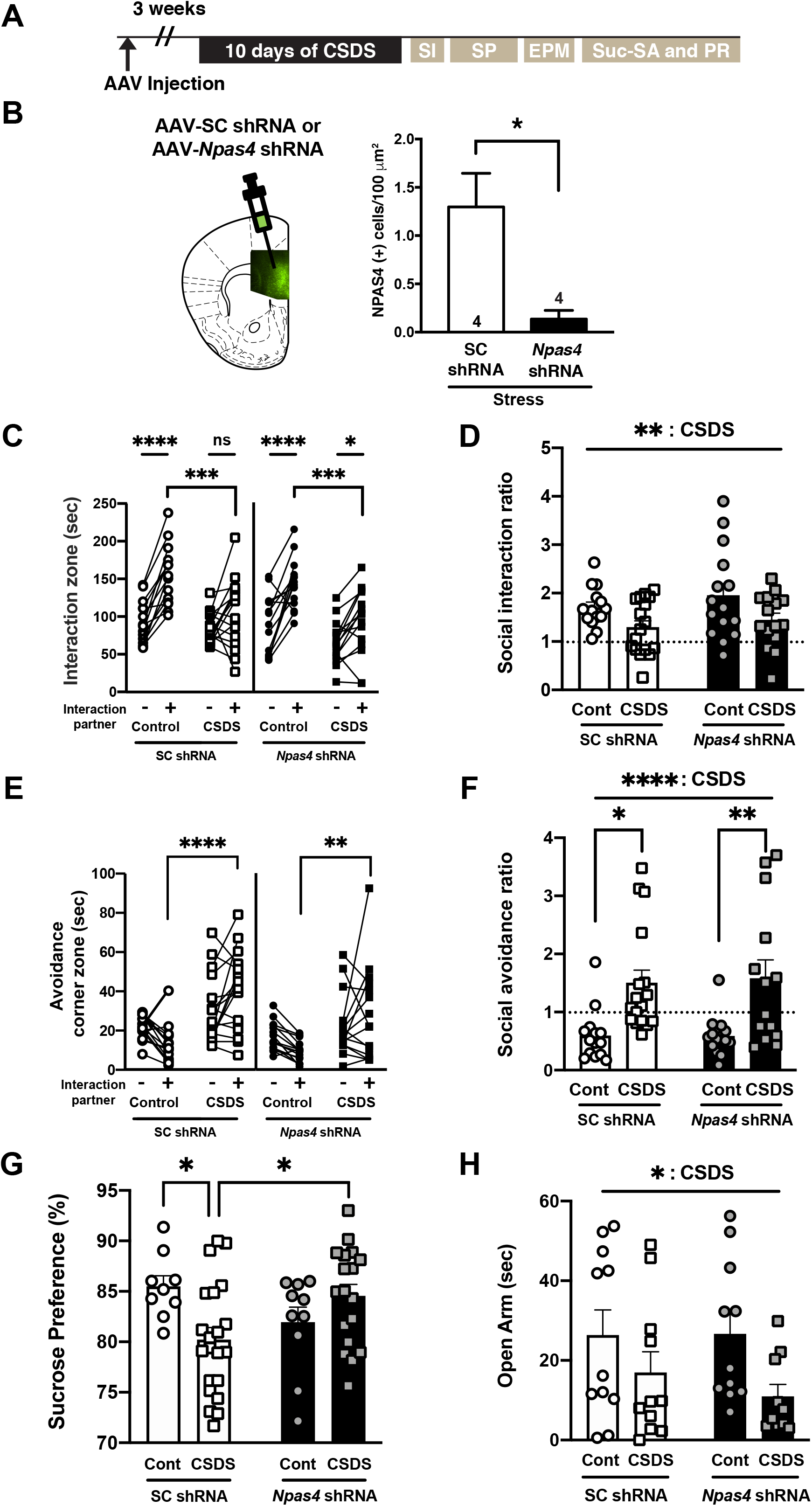
NPAS4 in the mPFC is required for CSDS-induced anhedonia-like behavior. (A) Schematic illustration of experimental timeline of behavioral test battery consisting of CSDS followed by social interaction (SI; C-F), sucrose preference (SP; G), elevated plus maze (EPM; H), sucrose self-administration, and progressive ratio testing (Suc-SA and PR; Figure 3 A-D). (B) AAV2-*Npas4* shRNA in the adult male mPFC decreases stress-induced NPAS4 protein expression. Left: representative image showing AAV2-shRNA expression viral vector-mediated eGFP expression in the adult mice mPFC. Right: quantification of NPAS4-positive cells/100 μm^2^ (n= 4/condition). (C) and (D) CSDS decreases the time spent in the social interaction zone (C) and the social interaction ratio (D) in SC shRNA^PFC^ and *Npas4* shRNA^PFC^ mice after CSDS (n=14-17/condition). (E) and (F) CSDS increases the time spent in the avoidance ccorner zone and social avoidance ratio in SC shRNA^PFC^ and *Npas4* shRNA^PFC^ mice (n=14-17/condition). (G) CSDS-induced reduction of sucrose preference is blocked by *Npas4* shRNA in the mPFC (F; n=9-21). (H) CSDS reduces time spent in open arms (sec) in SC shRNA^PFC^ and *Npas4* shRNA^PFC^ mice (n= 10-11).

Individuals who suffer from pathological stress often exhibit deficits in motivated, effort-based decision-making ^40-44^, though clinical studies have shown that some individuals can exhibit positive behavioral outcomes following stress (*i*.*e*. stress resilience) ^45^. To examine the role of NPAS4 in CSDS-induced changes in reward motivation, *Npas4* shRNA^PFC^ or SC shRNA^PFC^ mice were subjected to CSDS or the “no stress” condition, and then they were allowed to self-administer sucrose (sucrose SA) under operant conditions. After stable sucrose SA was established, we examined motivation to work for a sucrose reward using the progressive ratio (PR) schedule of reinforcement. Compared to SC shRNA^PFC^ controls, *Npas4* shRNA^PFC^ mice displayed no differences in acquisition of sucrose SA (Figure 3A) or operant discrimination learning (nosepokes in the active vs. inactive port) (Figure 3B). Interestingly, there was a statistical trend for *Npas4* shRNA^PFC^ to increase the PR breakpoint – the maximum number of nosepokes an animal was willing to perform to receive a single sucrose reward (Figure 3C), suggesting that reducing levels of mPFC NPAS4 might enhance reward motivation. Interestingly, when we compared SI and PR breakpoint (reward motivation) in CSDS-treated mice, we noted that SC shRNA^PFC^ control mice showed a strong positive correlation (Figure 3D). In other words, stress-susceptible mice that developed a decrease in social interaction following CSDS showed lower reward motivation; whereas, stress-resilient mice that developed an increase in SI after CSDS showed a high level of reward motivation, similar to human patients who present with beneficial psychological outcomes following stressful experiences ^45^. In stark contrast, mice lacking *Npas4* in the mPFC did not show a correlation of SI with motivation. Additionally, there was a significant difference in the slopes of correlation between SC shRNA^PFC^ and *Npas4* shRNA^PFC^ mice, suggesting that CSDS-induced mPFC NPAS4 influences natural reward motivation.

**Figure 3.**
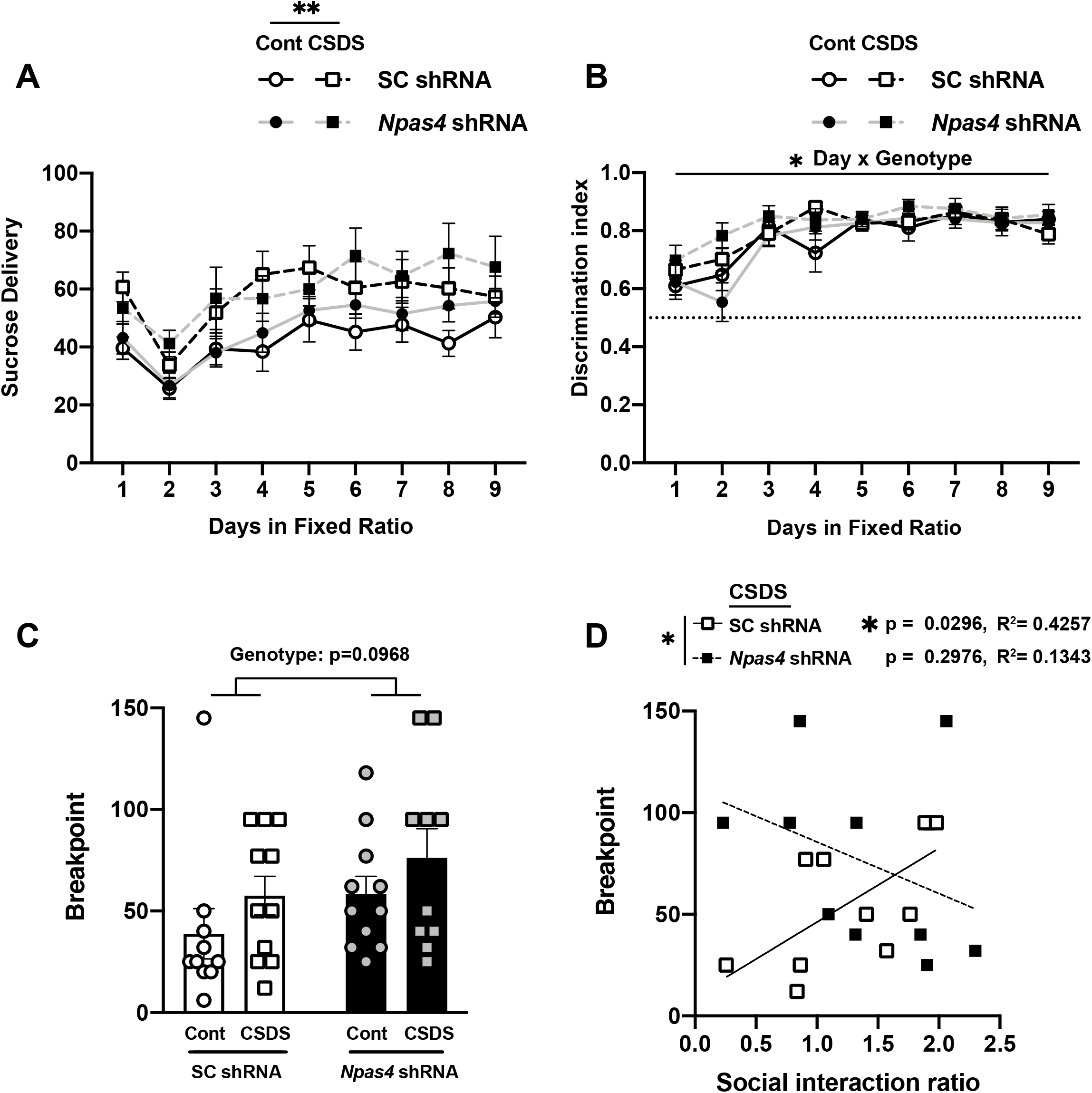
NPAS4 in the mPFC regulates effort-based motivated behavior during sucrose SA following CSDS. (A) and (B) Data plots showing the acquisition period of sucrose self-administration in SC shRNA^PFC^ and *Npas4* shRNA^PFC^ mice after CSDS or no stress control condition, with increased number of sucrose pellets received following CSDS (A) and no change in the discrimination ratio between the active and inactive nosepokes (B; n=10-11/group). (C) Data plot showing the maximum number of active nose pokes required to receive a sucrose reward (breakpoint) after SDS in the PR test of both SC shRNA^PFC^ and *Npas4* shRNA^PFC^ mice. *Npas4* shRNA^PFC^ mice demonstrated a trend for elevated PR breakpoint compared to control SC shRNA^PFC^ mice (n=10-11/group). (D) SC shRNA^PFC^ mice, but not *Npas4* shRNA^PFC^ mice, after CSDS exhibited significant positive correlations between the breakpoint and social interaction ratio.

### NPAS4 regulates CSDS-induced reductions in mPFC dendritic spine density

CSDS-induced reduction of dendritic spine density on mPFC pyramidal neurons is a putative pathophysiological underpinning of depression-associated behavior ^46-53^. As such, we quantified dendritic spine density on deep-layer pyramidal neurons in SC shRNA^PFC^ or *Npas4* shRNA^PFC^ mice after CSDS, compared to non-stressed mice. As expected, we observed a CSDS-induced reduction in dendritic spine density in SC shRNA control mice (Fig. 4A, top; 4B, left). In contrast, we observed no CSDS-induced changes in mPFC dendritic spine density in *Npas4* shRNA^PFC^ mice (Figure 4A, bottom; 4B, right), suggesting that NPAS4, directly or indirectly, is required for this chronic stress-induced structural synaptic change in the mPFC. Of note, no changes in mPFC dendritic spine density were observed in non-stressed *Npas4* shRNA^PFC^ mice (Figure 4A-B), indicating that steady-state dendritic spine density in adult mPFC pyramidal neurons of non-stressed animals does not require normal NPAS4 expression levels. In addition, neither *Npas4* shRNA nor CSDS produced any detectible changes in mean dendritic spine head diameter (Supplemental Figure S4). These data suggest that CSDS-induced NPAS4 is required for changes in spine density and that this change likely contributes to the regulation of anhedonia-like behavioral output following chronic stress.

**Figure 4.**
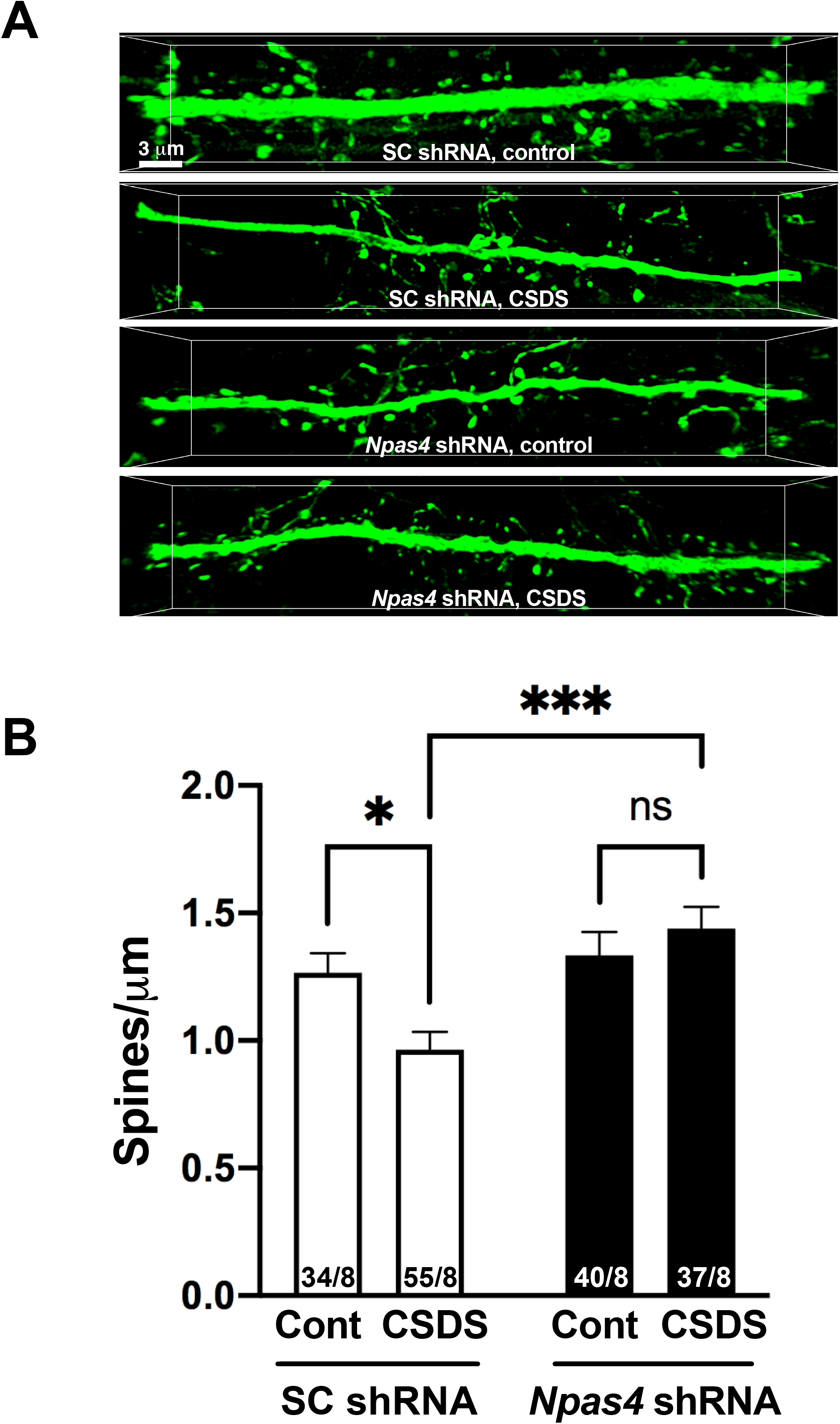
NPAS4 regulates CSDS-induced reductions in mPFC dendritic spine density. (A) and (B) NPAS4 regulates CSDS-induced reduction of dendritic spine density in the mPFC. (A) Representative images showing AAV2-shRNA expression viral vector-mediated eGFP expression. Scale bar, 3 μm. (B) Quantification of dendritic spine density of deep layer mPFC pyramidal neurons from SC shRNA^PFC^ and *Npas4* shRNA^PFC^ mice after CSDS or in no stress controls. (n= 34-55 branch/8 animals/condition). Data shown are mean ± SEM; *p < 0.05, ***p < 0.001. Also see Table S1 for detailed statistical analyses.

### NPAS4 regulates the expression of ribosomal and glutamatergic synapse genes

To analyze the influence of NPAS4 on the mPFC transcriptome, we performed RNA-seq analyses with mPFC tissue isolated from SC shRNA^PFC^ and *Npas4* shRNA^PFC^ mice. Of the ∼700 differentially expressed genes (DEGs, FDR <0.05, log_2_ (FC) > |0.3|) following *Npas4* mRNA knockdown in mPFC, 267 were down-regulated and 365 were up-regulated (Supplemental Table S2). Downregulated genes included *Gm1110, RP23-303F24*.*3, Spata3, Gm9843, and Defb1* and upregulated genes included *RP23-304I1*.*3, Gm20939, Arc, Igfn1*, and *A730015C16Rik* (Figure 5A, 5B). A subset of these DEGs was independently validated by qRT-PCR using independent mPFC samples isolated from SC shRNA^PFC^ and *Npas4* shRNA^PFC^ mice, including *Nfix* (Nuclear Factor I X), *Sst* (Somatostatin), *Nrp1* (Neuropilin 1), *Dhcr7* (7-Dehydrocholesterole Reductase), and *Ache* (Acetylcholinesterase) (Supplemental Figure S5). *Npas4* shRNA DEGs were significantly enriched in two PsychENCODE modules (Figure 5C) ^54, 55^; *Npas4* shRNA-downregulated DEGs showed significant PsychENCODE enrichment within gene module M15, an excitatory neuron module of genes that are associated with ribosome function and upregulated in Autism Spectrum Disorder (ASD) and Bipolar Disorder (BD), while *Npas4* shRNA-upregulated DEGs showed significant enrichment in gene module M1, an excitatory neuron module of genes that are associated with glutamate-driven excitability of neurons that have been shown to be downregulated in ASD ^54^. Additionally, pathway analyses of *Npas4* shRNA downregulated DEGs revealed significant enrichment of genes linked to ribosome function and normal translation mechanisms, and upregulated DEGs showed significant enrichment of glutamatergic synapse-related genes that are important for synaptic signaling and organization (Figure 5D). We detected 92 total ribosome-related genes in the mPFC that are classified in the pathway “co-translational protein targeting to membrane”, and of these, 68 were significantly downregulated and 2 upregulated (p < 0.05) by *Npas4* shRNA (Figure 4E). Interestingly, RNA-seq from human postmortem brains (BA8/9) of male MDD patients indicated significant enrichment of differentially expressed genes (p < 0.05) in ribosome-related pathways, identifying 57 significantly upregulated genes in MDD patients (Figure 5E) ^39^. Moreover, the majority (66.2%) of the *Npas4* shRNA-mediated significantly downregulated genes from our analysis overlapped with upregulated genes in human MDD patients (Figure 5E). Finally, ChIP-seq with NPAS4 antibody in hippocampal neurons previously demonstrated genome-wide DNA binding of NPAS4 ^34^; NPAS4 directly associates with around 55% (42 of 77) of ribosome-related genes classified in the pathway “co-translational protein targeting membrane” (Figure 5F). Together, these data suggest that NPAS4 in the mPFC regulates numerous genes related to glutamatergic synapse regulation and normal ribosomal function, which potentially contributes to mPFC hypofrontality and stress-related anhedonia. Moreover, the enrichment of NPAS4-regulated genes in multiple neuropsychiatric disorders, including MDD ^39^, positions *Npas4* as an important transcription factor in brain health and disease, opening a future research pathway for the study of how these DEGs contribute to depression and anhedonia-related behaviors in both preclinical and clinical models.

**Figure 5.**
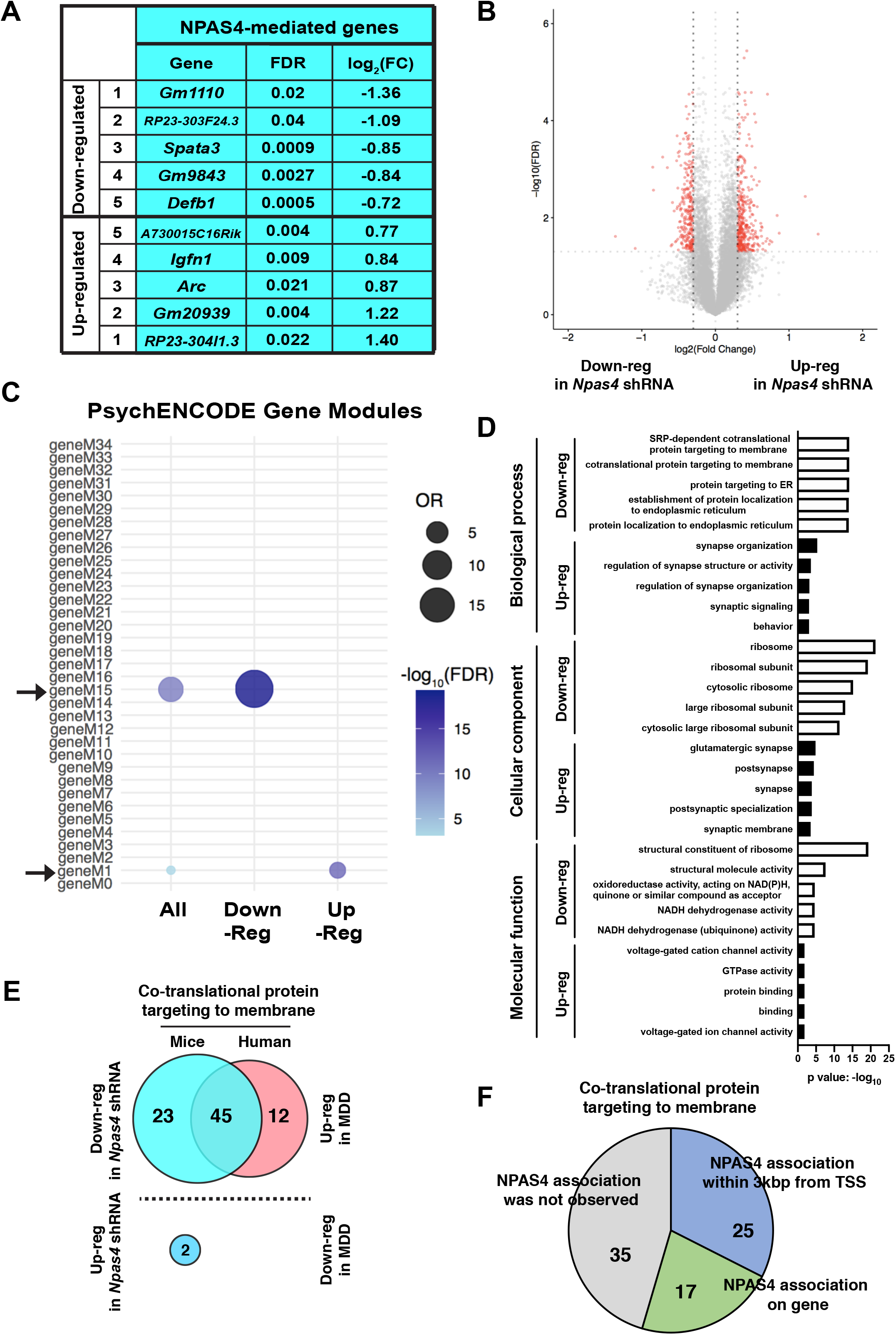
NPAS4 regulates the expression of ribosomal and glutamatergic synapse genes. (A) and (B) List of top differentially expressed genes in mPFC of *Npas4* shRNA^PFC^ mice (A) and corresponding volcano plot of all significant DEGs (FDR <0.05, log_2_ (FC) > |0.3|, red) compared to those that were not significant (gray; B). (C) *Npas4* DEG enrichment in gene modules that are dysregulated in neuropsychiatric disorders; Modules M1 and M15, as shown by PsychENCODE. (D) Gene ontology analysis of down- and up-regulated DEGs in *Npas4* shRNA^PFC^ mice. (E) Overlap of significantly differential expression genes (p < 0.05) in *Npas4* shRNA^PFC^ mice (left; blue) and differential expression genes (p < 0.05) in BA8/9 of human MDD patients (right; pink). (F) ChIP-seq analysis of NPAS4 association with significant ribosome-related differential expression genes identified from this study.

## Discussion

Here we find that social defeat stress (acute or chronic) induces rapid and transient expression of NPAS4 in mPFC neurons, and that NPAS4 in the mPFC is required for CSDS-induced anhedonia-like behavior, changes in effort-based reward seeking-motivated behavior, and CSDS-induced dendritic spine loss on mPFC pyramidal neurons. However, mPFC NPAS4 was not required for CSDS-induced social avoidance or anxiety-like behavior, suggesting that CSDS produces different depressive-like phenotypes through distinct molecular and/or circuit mechanisms. Finally, we discovered that *Npas4* shRNA dysregulated ∼632 mPFC genes, including upregulation of genes linked to glutamatergic synapses, suggesting that following CSDS, NPAS4 could directly or indirectly downregulate these synapse-related genes to enable structural dendritic spine reduction. We also detected strong enrichment of downregulated ribosomal genes, many of which are also dysregulated in human MDD, providing evidence that some of these genes could be potential biomarkers for depression. Together, our findings reveal a novel and essential role for NPAS4 in transcriptomic regulation of mPFC excitatory neurons that precede the induction of anhedonia-related behavior and structural synaptic changes produced by CSDS.

NPAS4 is a neuronal-specific, synaptic activity-regulated and experience-dependent transcription factor that regulates excitatory/inhibitory synapse balance and synaptic transmission ^28, 29, 31, 32, 34, 56^, promoting cell type-specific gene programs and cellular responses. Synaptic activity-dependent induction of NPAS4 in pyramidal neurons reduces excitatory synaptic transmission onto these neurons ^29^ and decreases excitatory synaptic inputs ^30^, consistent with our finding that NPAS4 is required for CSDS-induced loss of mPFC pyramidal neuron dendritic spine density. While one report indicated that CSDS-induced reduction of dendritic spine density is associated with social avoidance phenotypes ^53^, we observed that mPFC NPAS4 reduction selectively blocked CSDS-induced spine loss and anhedonia, but not social avoidance and anxiety-like behavior, suggesting that deep-layer mPFC pyramidal cell spine loss, *per se*, is not required for the expression of social- and anxiety-related phenotypes in the CSDS model. Although we were unable to study females in our CSDS model, chronic exposure to stress hormones, chronic mild unpredictable stress, and chronic restraint stress all induce anhedonia-like behavior and dendritic spine loss in both sexes ^20, 48, 51, 57-64^. Moreover, PFC pyramidal cell dendritic spine density is also reduced in human postmortem brains of individuals diagnosed with anhedonia-associated neuropsychiatric disorders, such as SCZ, BD, and MDD ^51,57, 65-72^. These data support the possible functional relationship between PFC spine density and anhedonia, analogous to what we see in mice after *Npas4* manipulation in this brain region.

NPAS4 in cultured neurons regulates a large, cell type-specific program of gene expression, including key targets like brain-derived neurotrophic factor (BDNF), that alter E/I synapse balance ^10, 28-32, 36^. However, social defeat stress failed to induce *Bdnf* mRNA in mPFC and *Npas4* knockdown did not alter basal mPFC *Bdnf* expression (data not shown and Supplemental Table S2), suggesting that *Bdnf* is not a key downstream target of mPFC NPAS4 in the context of CSDS. Our RNA-seq analysis of mPFC tissues, with or without *Npas4* shRNA, revealed an abundance of significant DEGs (Figure 5). Of the upregulated DEGs, gene ontology (GO) pathway analysis revealed enrichment of genes linked to glutamatergic synaptic transmission and excitability, and PsychENCODE analysis identified a neuronal module of genes linked to glutamatergic excitability that are downregulated in Autism Spectrum Disorders ^54^, suggesting the possibility that CSDS-induced mPFC dendritic spine density loss is produced, in part, by one or more of these synapse-linked genes that are downregulated following stress-induced mPFC NPAS4 expression. Interestingly, 22% of these upregulated DEGs overlapped with NPAS4 target genes identified by ChIP-seq analysis from cultured pyramidal neurons ^73^, suggesting that some of the upregulated, synapse-related DEGs could be direct NPAS4 gene targets. In contrast to the upregulated genes, *Npas4* shRNA-downregulated genes showed strong enrichment for ribosomal function and a PsychENCODE module (M15) of excitatory neuron genes associated with ribosome function that is upregulated in ASD and BD, and more than half of these downregulated *Npas4* shRNA genes are associated with NPAS4 protein (Figure 5F). While the functional relevance of ribosome gene enrichment is unclear, the marked enrichment of ribosome-related DEGs is very striking. Microarray analysis of blood samples from stress-vulnerable vs. stress-resistant adults found that DEGs were most markedly enriched in ribosome-related pathways and were upregulated based on stress vulnerability ^74^. Additionally, RNA-seq analyses from orbitofrontal cortex of postmortem human brains with SCZ, BD, and MDD also identified DEGs enriched for the ribosomal pathway, most of which were upregulated in patient samples ^75^. Together, these data suggest that NPAS4 regulates the expression of numerous mPFC genes, including those linked to ribosome function and glutamatergic synapses, as well as genes dysregulated in patients with anhedonia-associated neuropsychiatric disorders, and that NPAS4 might promote anhedonia and hypofrontality through one or more of these gene targets.

Overall, our findings reveal a novel role for mPFC NPAS4 in CSDS-induced anhedonia and pyramidal neuron dendritic spine loss, but not in the development of social avoidance or anxiety-like behavior. We found that mPFC NPAS4 regulates hundreds of genes, including clusters of genes linked to glutamatergic synapse function and ribosomal function, both of which are well-positioned to alter neuronal function. Therapeutic strategies targeting *Npas4*-regulated pathways could be a novel approach to develop new treatments for hypofrontality and anhedonia-related symptoms in patients struggling with depression, bipolar disorder, and other stress-related neuropsychiatric disorders.

## Acknowledgments

The authors thank Rachel Penrod, Laura Smith, Yuhong Guo, Ben Zirlin, and Sara Pilling for technical assistance, comments on the manuscript, and helpful discussions. B.W.H. was supported by an NIH predoctoral fellowship (F31 DA048557 and T32 DA07288). B.M.S. was supported by an NIH postdoctoral fellowship (F32 DA050427). M.T. was supported by a NARSAD Young Investigator Award from the Brain & Behavior Research Foundation (Grant #22765). This work was supported by grants from the NIH (UL1 TR001450 to M.T., R01 DA032708 and DA046373 to C.W.C.)

