## Supplemental file and figures for "NPAS4 in the medial prefrontal cortex mediates chronic social defeat stress-induced anhedonia and dendritic spine loss": Hughes_Supplemental figure and legends_20210902.pdf

**Inventory of supplemental information**

Supplemental Methods and Materials

References

Supplemental figures

Figure S1. Related to Figure 1

Figure S2. Related to Figure 2

Figure S3. Related to Figure 4

Figure S4. Related to Figure 5

Supplemental Tables

Table S1. Detailed statistics information related to Figure 1-5 and Supplemental Figure S1-S4

Table S2. RNA-seq data Related to Figure 4

Table S3. Primers used in gene expression analyses

### Supplemental Methods and Materials

#### Animals

C57BL/6 adult male mice were purchased from Jackson Laboratory (ME) and tested between 8-20 weeks of age. Mice were allowed access to food and water *ad libitum* and were kept on a 12 hr light-dark cycle. All procedures were in accordance with Institutional Animal Care and Use (IACUC) guidelines.

#### qRT-PCR for gene expression

Brain tissue was collected at the described time point after social defeat stress, or no stress control condition, and kept frozen at -80°C until processed for the following steps. RNA isolation, reverse transcription, and quantitative real-time PCR were carried out as described previously [1]. Mouse tissue samples were homogenized in QIAzol solution and processed for RNA purification using the miRNeasy kit (QIAGEN, MD) following the manufacture's protocol. The total RNA was reverse transcribed using SuperScript III (Invitrogen) with a random hexamer primer following the manufacturer's instructions. Quantitative PCR (qPCR) was performed using SYBR Green (Bio-Rad, CA). The level of mRNA expression was analyzed by the fold change relative to *Gapdh* or *$\beta$-tubulin* expression. The relative mRNA level was analyzed as the difference from experimental condition relative to controls. See the supplemental Table S3 for primer sequences.

#### Immunohistochemistry

Mouse brains were fixed overnight in 4% PFA in 1x PBS and transferred to a 30% sucrose solution in 1x PBS before slicing (40 or 50 μm) with a microtome. The slices were permeabilized and blocked in 3% BSA, 0.3% Triton X-100, 0.2% Tween-20, 3% normal donkey or goat serum in PBS, then incubated with primary antibodies: anti-GFP (1:1000, Aves Labs, Inc, OR; 1:1000-10,000, Invitrogen), anti-NPAS4 (1:1000, rabbit, kindly provided by Dr. Michael Greenberg's lab), anti-CaMKIIα (1:1000, Enzo, NY, 6G9), anti-somatostatin(SST) (1:1000, Millipore MAB354), and anti-Parbalbumin(PV) (1:1000, Millipore MAB1572) in blocking buffer at room temperature for 2-4 hours or at 4°C overnight. Following a series of 1x PBS rinses, slices were incubated for 1-3 hr at room temperature with secondary antibodies (Donkey anti-rabbit 488, Goat anti-mouse 594, Donkey anti-mouse Cy3, or Donkey anti-chicken 488) while protected from light. Slices were counterstained with Hoechst, mounted and coverslipped on glass slides using AquaMount (Thermo scientific, MA) or ProlongGold (Thermo scientific, MA) and analyzed with confocal microscopy (Zeiss LSM 880). The expression level of NPAS4 protein in each cell was measured using ImageJ software in CaMKIIα-positive cells under experimenter-blinded conditions.

**Social interaction assay**

Social Interaction (SI) assay was performed as previously described [2, 3]. The social interaction assay was performed 24 hours after the last CSDS procedure. The assay was performed in an open field arena (44 cm x 44 cm) with the social target's holding cage. The time mice spent in the interaction zone (8 cm from the social target) was examined for 5 min in the absence and then the presence of a novel CD1 mouse, of which the experimental animal never met, under dim red light using AnyMaze 5.1 (Stoelting Co, Wood Dale) or Ethovision 3.0 software (Noldus, Leesburg, VA). The social interaction ratio was calculated as the time spent in the social interaction zone in the presence of an interaction partner divided by the time in the absence.

**Sucrose preference test**

The sucrose preference procedure was performed as previously described [4]. Single-housed mice were provided with Division of Laboratory Animal Resources-approved tap water in 2 identical double-ball-bearing sipper-style bottles for 2 days, followed by 2 days of 1% (w/v) sucrose solution in tap water to allow for acclimation. Mice were then given 1 bottle containing tap water and another containing the 1% sucrose solution. Consumption from each bottle was measured every 24 hours for 4 days, and bottle positions were swapped each day to avoid potential side bias. The sucrose preference was calculated by percentage of 1% sucrose consumption volume divided by total liquid consumption volume (sucrose + tap water). Measurement of liquid consumption volume was performed with experimenters blinded to conditions.

**Elevated plus maze**

The elevated plus maze (EPM) was conducted under the bright light (80 Lux at the closed arm) as previously performed [1]. Mice were positioned in the center of the maze, and behavior was recorded by video tracking using AnyMaze 5.1 or Ethovision 3.0 software as previously performed [5]. The time spent in the open arms was recorded for 5 min.

**Sucrose self-administration assay**

Sucrose self-administration was conducted for 2 hr at the same time each day for 9-10 days of acquisition training, followed by a progressive ratio schedule as described previously[1]. Briefly, sucrose availability was signaled both by the house light and a light above the active nosepoke hole. Following a poke in the active hole, both availability lights went off and a cue light inside the nose poke hole was illuminated. Sucrose pellets were delivered immediately upon the active nosepoke, followed by a 10 s time-out period. Nose pokes in the inactive hole were without programmed consequences. During a progressive ratio schedule of reinforcement, the requirements for a sucrose delivery were increased on a subsequent sucrose delivery in an exponential manner. Animals were allowed to self-administer until they failed to earn a sucrose pellet in a 60 min time frame. The last sucrose pellet achieved is reported as an indicator of how much animals consumed before reaching breakpoint.

**Dendritic spine morphometric analyses**

Mouse brains were collected with rapid live decapitation 24 hr after the social interaction assay and fixed overnight in 4% PFA in 1x PB, then transferred to a 30% sucrose solution in 1x PB before slicing (40 μm) with a vibratome. Deep layer eGFP-expressing pyramidal neurons in the prelimbic cortex were sampled for dendritic spine analyses as described previously [6]. Briefly, proximal apical dendrites were imaged with a Leica SP8 laser scanning confocal microscope equipped with HyD detectors for enhanced sensitivity. Dendritic spine segments were selected only if they satisfied the following criteria: 1) could clearly be traced back to a cell body of origin, 2) were not obfuscated by other dendrites, and 3) were proximal to the branch point separating the apical tuft from the proximal apical dendrite. Images were collected with a 63X oil immersion objective (1.4 N.A.) at 1024x1024 frame size, 4.1X digital zoom, and a 0.1μm Z-step size

(0.04x0.04x0.1 μm voxel size). Pinhole was set at 0.8 airy units and held constant. Laser power and gain were empirically determined and then held relatively constant, only adjusting to avoid saturated voxels. Huygens Software (Scientific Volume Imaging, Hilversum NL) was used to deconvolve 3D Z-stacks. Deconvolved Z-stacks were then imported into Imaris (version 9.0.1) software (Bitplane, Zurich CH). The filament tool was then used to trace and assign the dendrite shaft. Dendritic spines were then semi-automatically traced using the autopath function, and an automatic threshold was used to determine dendritic spine head diameter. Variables exported included the average spine head diameter (in μm) as well as the number of dendritic spines per μm of dendrite (spine density). 3-10 segments were sampled per animal, and the average spine head diameter and the spine density were calculated for each segment. Data for each variable was then expressed as number of spine segments/number of animals. All analyses were performed under experimenter-blinded conditions.

**RNA-seq and bioinformatic analysis**

Total RNA was isolated from AAV2-mediated eGFP-positive mPFC slices using the QIAGEN RNA purification kit, as described above. Sequencing was performed by BGI genomics using PolyA mRNA isolation, directional RNA-seq library preparation, and a BIGSeq-500 sequencer. Reads were aligned to the mouse mm10 reference genome using STAR (v2.7.1a) [7]. Only uniquely-mapped reads were retained for further analyses. Quality control metrics were assessed by Picard tool (http://broadinstitute.github.io/picard/). Gencode annotation for mm10 (version M21) was used as reference alignment annotation and downstream quantification. Gene level expression was calculated using HTseq (v0.9.1) [8] using the intersection-strict mode by exon. Counts were calculated based on protein-coding genes from the annotation file.

#### Differential gene expression

Counts were normalized using counts per million reads (CPM). Genes with no reads were removed. Differential expression analysis was performed in R using linear modeling as following: lm(gene expression ~ Treatment + Batch). We estimated log2 fold changes and P-values. P-values were adjusted for multiple comparisons using a Benjamini-Hochberg correction (FDR). Differentially expressed genes were analyzed at FDR<0.05. Mouse Gene IDs were translated into Human Gene IDs using the biomaRt package (v2.46.0) in R [9].

#### Gene ontology analyses

The functional annotation of differentially expressed and co-expressed genes was performed using GOstats [10]. A Benjamini-Hochberg FDR (FDR<0.05) was applied as a multiple comparison adjustment.

#### Gene set enrichment

Gene set enrichment was performed in R using Fisher's exact test with the following parameters: alternative = "greater", confidence level = 0.95. We reported Odds Ratio (OR) and Benjamini-Hochberg adjusted P-values (FDR).

#### Statistics

One-way, two-way, and three-way analyses of variance (ANOVAs) with or without repeated-measures (RM) were used, followed by Bonferroni or Tukey post hoc tests when a significant interaction was revealed, to analyze mRNA expression, number of NPAS4 (+) cells, NPAS4 protein expression in each cell, percentage of CaMKIIα(+) cells, social interaction, social aversion, sucrose preference, elevated plus maze, sucrose self-administration acquisition and discrimination, dendritic spine morphometric data, and breakpoint in the progressive ratio test. All statistics were performed using GraphPad Prism, except SPSS software was used to handle complex datasets (e.g., three-way ANOVAs). Statistical outliers were detected using a Grubbs test and excluded from analysis. All data are presented as the mean ± SEM. Significance was shown as * = $p < 0.05$, ** = $p < 0.01$, *** = $p < 0.001$, **** = $p < 0.0001$, and non-significant values were either not noted or shown as n.s.

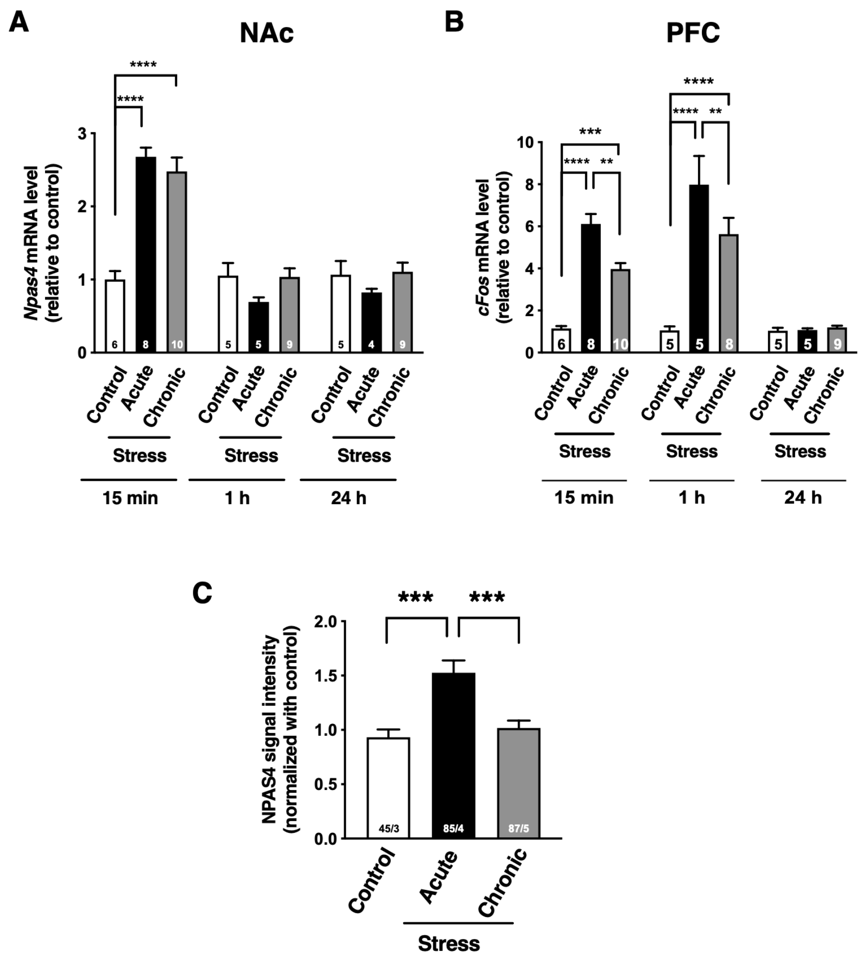


**Supplemental Figure S1. Social defeat stress induces NPAS4 and cFos expression in the NAc and mPFC**

(A) Quantification of *Npas4* mRNA expression in the NAc following acute and chronic social defeat stress at 15 min, 1 hr, and 24 hr. (n= 5-10/condition).

(B) Quantification of *cFos* mRNA expression in the mPFC following acute and chronic social defeat stress at 15 min, 1hr, and 24 hr. (n= 5-10/condition).

(C) Data plot represents fold change of NPAS4 signal intensity in CaMKII$\alpha$-positive pyramidal excitatory neurons of the mPFC (n= 45-87 cells/3-5 animals/condition).

Data shown are mean ± SEM; $^{**}p < 0.01$, $^{***}p < 0.001$, $^{****}p < 0.0001$. Also see Table S1 for detailed statistical analyses.

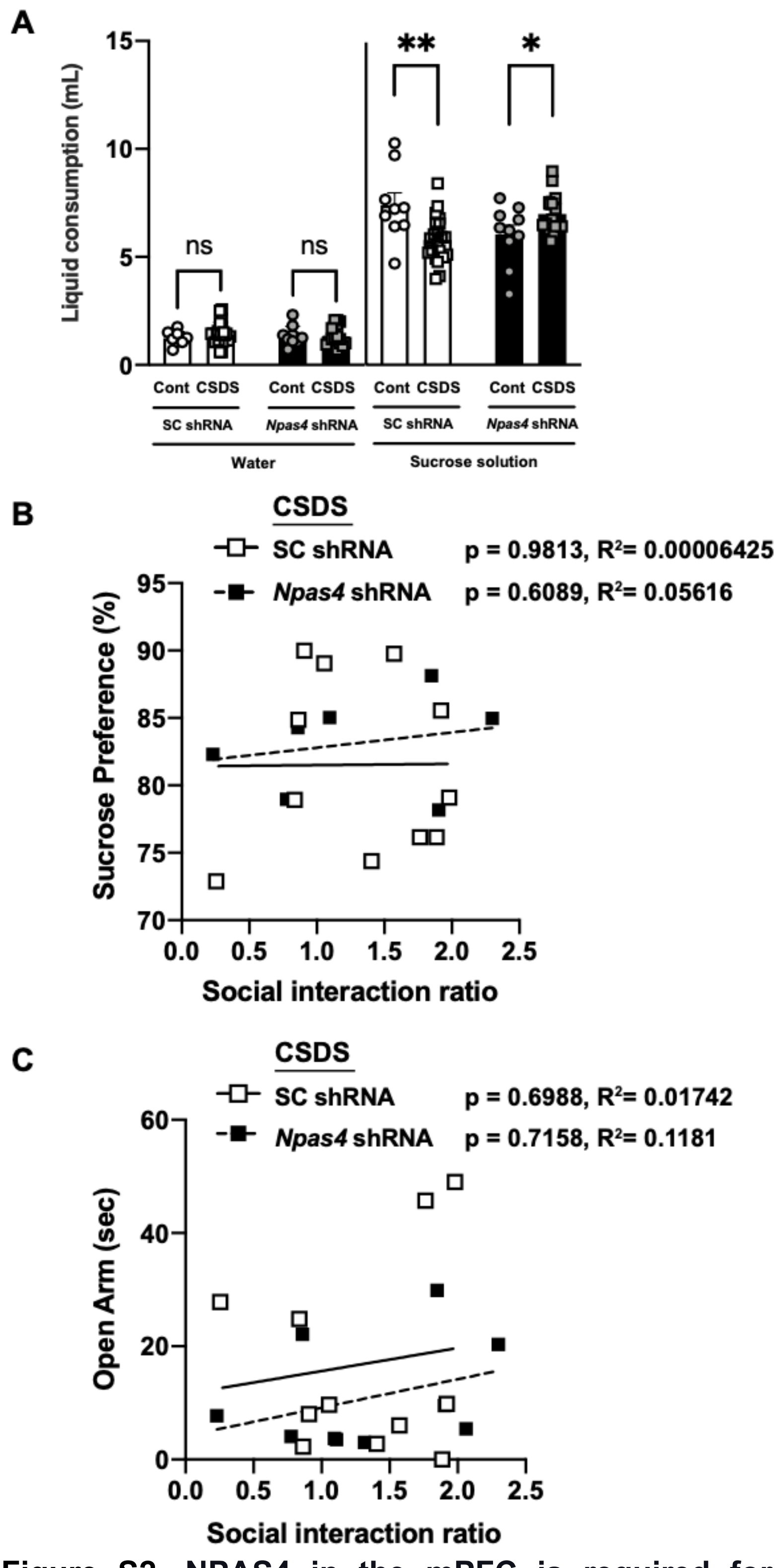


**Supplemental Figure S2. NPAS4 in the mPFC is required for CSDS-induced anhedonia-like behavior that is not correlated with social interaction ratio**

(A) Chart represents the amount of consumption of water and 1% sucrose solution in each day SC shRNA$^{PFC}$ and *Npas4* shRNA$^{PFC}$ mice after CSDS. (n= 9-21/condition).

(B) CSDS-induced reduction of sucrose preference was not correlated with CSDS-induced changes in social interaction ratios in SC shRNA$^{PFC}$ and *Npas4* shRNA$^{PFC}$ mice. (n=7-11/condition)

(B) CSDS-induced reduction of the time in the open arm (sec) was not correlated with CSDS-induced changes in social interaction ratios in SC shRNA$^{PFC}$ and *Npas4* shRNA$^{PFC}$ mice. (n=10-11/condition)

Data shown are mean ± SEM; *$p < 0.05$, **$p < 0.01$ Also see Table S1 for detailed statistical analyses.

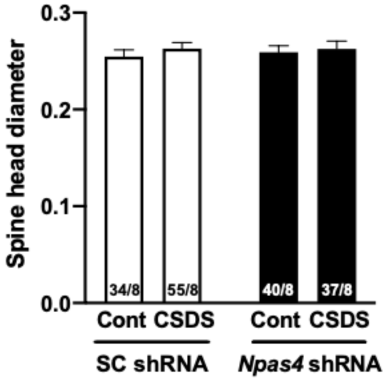


**Supplemental Figure S3. mPFC dendritic spine morphological analyses in the mPFC of SC shRNA$^{PFC}$ and *Npas4* shRNA$^{PFC}$ mice after CSDS**

Data plots represent spine head diameter of AAV2-SC shRNA or *Npas4* shRNA viral vector-mediated eGFP-positive mPFC pyramidal neurons after CSDS. (n= 34-55 branch/8 animals/condition).

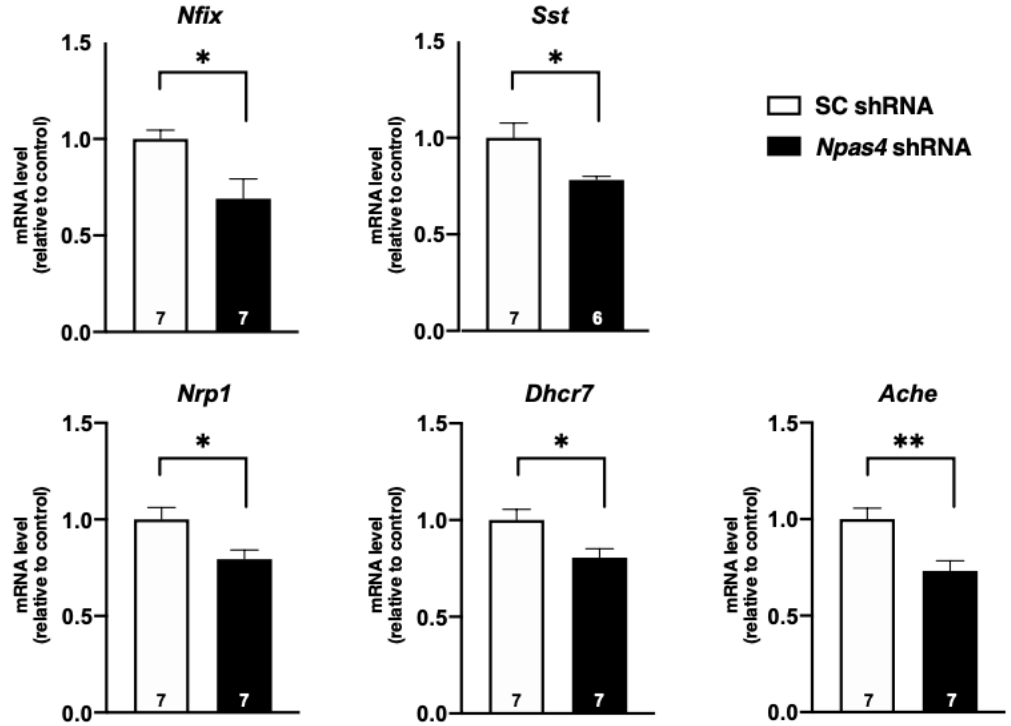


**Supplemental Figure S4. Differential expression genes in the mPFC of *Npas4* shRNA mice**

Data plots represent the relative mRNA expression in the mPFC of SC shRNA and *Npas4* shRNA mice. (n= 6-7/condition).

Data shown are mean ± SEM; *p < 0.05, **p < 0.01. Also see Table S1 for detailed statistical analyses.
